# Inferring the distributions of fitness effects and proportions of strongly deleterious mutations

**DOI:** 10.1101/2022.11.16.516724

**Authors:** Anders P. Charmouh, Greta Bocedi, Matthew Hartfield

## Abstract

The distribution of fitness effects is a key property in evolutionary genetics as it has implications for several evolutionary phenomena including the evolution of sex and mating systems, the rate of adaptive evolution, and the prevalence of deleterious mutations. Despite the distribution of fitness effects being extensively studied, the effects of strongly deleterious mutations are difficult to infer since such mutations are unlikely to be present in samples of haplotypes, so genetic data may contain very little information about them. Recent work has attempted to correct for this issue by expanding the classic gamma-distributed model to explicitly account for strongly deleterious mutations. Here, we use simulations to investigate one such method, adding a parameter (*p_lth_*) to capture the proportion of strongly deleterious mutations. We show that *p_lth_* can improve the model fit when applied to individual species but can underestimate the true proportion of strongly deleterious mutations. The parameter can also artificially maximize the likelihood when used to jointly infer a distribution of fitness effects from multiple species. As *p_lth_* and related parameters are used in current inference algorithms, our results are relevant with respect to avoiding model artifacts and improving future tools for inferring the distribution of fitness effects.

## Introduction

The distribution of fitness effects (DFE) among new mutations has long been recognized as having a fundamental importance in evolutionary genetics since the shape of this distribution affects many evolutionary phenomena. For example, the DFE governs the rate of adaptive evolution (Fisher 1930; Schultz and Lynch 1997; Spigler et al. 2017), the frequency (and hence prevalence) of mutations with different effect sizes (Haldane 1937; Agrawal and Whitlock 2012) and has important implication for phenomena such as inbreeding avoidance, the evolution of sex and mating system evolution (Kondrashov 1985; Hartfield and Keightley 2012; Hedrick and Garcia-Dorado 2016). Because of this, the DFE has been extensively studied over several decades (for a review, see Eyre-Walker and Keightley 2007). Many important results on the DFE have emerged from this large body of literature; the DFE differs among species (Charlesworth and Eyre-Walker 2006; Castellano et al. 2018) and the shape depends on how well-adapted the population is and its effective population size *N_e_* (Eyre-Walker 2002; Lynch and Conery 2003; Goldstein 2013; Huber et al. 2017). While the full DFE is probably complex and multi-modal, parts of the DFE of deleterious mutations can be modeled as a leptokurtic gamma distribution (Loewe and Charlesworth 2006; Silander et al. 2007) or by a series of discrete bins showing the proportions of mutations with different effects (Keightley and Eyre-Walker 2010).

Most of these results on the DFE have been obtained through two experimental techniques: mutation-inducing experiments and mutation accumulation experiments. In the former, mutations are induced and their effects are evaluated by comparing different strains of a model organism using some trait, such as fertility or growth rate, as a fitness proxy (Sanjuán et al. 2004). In the latter and more common approach, an ancestral strain is split into small, separate populations which are kept for many generations under favourable conditions that impose very little selection (Ohnishi 1977; Halligan and Keightley 2009; Böndel et al. 2019). The effects of accumulated mutations can then be evaluated by comparing the fitness proxy of the population strains with that of the ancestral strain. The DFE can also be estimated by sampling and comparing haplotypes from individuals in the wild (Dobzhansky and Wright 1941; Crow and Temin 1964; Eyre-Walker et al. 2006; Castellano et al. 2018). Data from the sampled haplotypes are often analysed in the form of a site frequency spectrum (SFS) which describes the frequency with which different alleles segregates in the sampled population (Fisher 1931; Wright 193 8; Evans et al. 2007).

One major challenge in DFE inference is measuring the occurrence of strongly deleterious mutations. Assuming deleterious alleles are not completely recessive, their mutation-selection frequency is generally proportional to 1/*s*, where *s* is the mutation’s selection coefficient (Wright 1937; Crow and Kimura 1970), and mutations are less likely to be present in a sample of haplotypes as they become more strongly deleterious due to the large selective disadvantage they entail. In other words, the probability distribution of mutational effects in segregating mutations only will almost certainly be different from the probability distribution of all mutations, since strongly deleterious mutations (with s» l/2*N_e_*) have a tiny probability of showing up in a population sample (Crow and Kimura 1970; Reich and Lander 2001; Rice et al. 2015). This emphasises the importance of distinguishing between an *observed* DFE, consisting of alleles segregating in the gene pool of a population at a given timepoint, and an *input* DFE that represents the actual probability distribution of mutational effects as they arise by mutation and are added to the gene pool.

Much work has been done on inferring what proportion of the DFE consists of strongly deleterious mutations; a study on human polymorphism data puts the estimate at <15% (Eyre-Walker et al. 2006). However, this proportion differs greatly among different branches of the tree of life. The proportion of lethal mutations was found to be approximately 40% in the vesicular stomatitis virus (Sanjuán et al. 2004), and even above 70% in *Drosophila* (Keightley and Eyre-Walker 2007). However, two mutations which are both described as strongly deleterious can still be orders of magnitude different in effect as mutations with, for example, *s*=10 ^2^/*N_e_* and *s*=10 ^5^/*N_e_* both satisfy *s* » 1/*2N_e_.* As a result, modem DFE inference algorithms such as DFE-alpha (Eyre-Walker and Keightley 2007; Keightley et al. 2016) and polyDFE (Tataru et al. 2017; Tataru and Bataillon 2020) which all work by analysing a SFS, may struggle to describe the extent to which many different strongly deleterious mutations arise. This is partly because very strongly deleterious mutations are unlikely to show up in a genome sample. However, the number of non­segregating sites in a genome sample (some of which are presumed to be under strong purifying selection) may provide information about the rate at which strongly deleterious mutations occur. In any case, it is hard to differentiate the effect sizes of mutations once the effect size is large, since such a mutation will have essentially no probability of appearing in a genome sample (Galtier and Rousselle 2020).

A few studies have attempted to account for these strongly deleterious mutations by fitting a DFE under the assumption that a certain fraction of mutations are so strongly deleterious that they will remain absent from a haplotype sample, regardless of *N_e_* (Eyre-Walker et al. 2006; Boyko et al. 2008; Elyashiv et al. 2010; Kim et al. 2017; Galtier and Rousselle 2020). In particular, Galtier and Rousselle (2020) used DFE methods to infer the mean *N_e_s* of mutations among species, thereby obtaining estimate of the difference in *N_e_* between different species under the assumption that species shared a common, gamma­distributed DFE. The field of DFE research has often parameterized gamma distributions through the usage of a shape parameter, often denoted *β*, and a mean value (Keightley and Eyre-Walker 2007). The optimal shape parameter of the gamma-distributed DFE varied greatly among species, but a good fit for similar shape parameters was obtained when a fraction of strongly selected mutations, with an effect size making them deleterious to the point of being “essentially independent of *N_e_*” was included in the SFSs. Here, *N_e_-* independent mutations were defined as those deleterious enough so that their probability of showing up in a haplotype sample is essentially zero regardless of *N_e_*. Galtier and Rousselle (2020) implemented this approach was implemented by assuming that a proportion of 0≤*p_lth_* ≤l mutations were either lethal or so strongly selected that they never appeared in a SFS; each entry in the selected SFS was subsequently scaled by 1–*p _lth_.* The method yielded a high model likelihood but produced very surprising results such that estimates of the proportion of strongly deleterious mutations (including lethal and nearly lethal mutations) in *Drosophila* being over 50%, which is well above an order of magnitude larger than previous estimates (Lethal or very strongly deleterious mutations are estimated to occur at a rate of approx. 10% of the rate of mildly deleterious mutations; Simmons and Crow 1977; Crow and Simmons 1983). While this method provides a way of accounting for strongly deleterious mutations that will be absent from genomic data, the properties and accuracies of this approach has yet to be fully investigated.

Here, we investigate to what extent explicitly accounting for strongly deleterious mutations in a gamma distributed DFE model improves the accuracy of inference when inferring an input DFE from simulated and genomic data. We do this by formulating a genetically explicit individual-based Wright-Fisher model to simulate steady-state observed DFEs under different strengths of selective effects, then infer the DFEs using the state-of-the-art DFE inference software polyDFE (Tataru et al. 2017). We show that explicitly accounting for a proportion of strongly deleterious mutations can increase the accuracy of inference when inferring the DFE of a single population. However, we also demonstrate that including this parameter artificially increases the inferred proportion of strongly deleterious mutations when considering SFSs from multiple different populations. We further show that the *p_lth_* resulting in the best model fit is not equivalent to the true proportion of lethal or strongly deleterious mutations in both cases.

## Methods

To simulate the observed DFE of a population, we model a diploid, sexually reproducing Wright-Fisher population with discrete generations and a default population size of *N* = 500. Individuals have two different chromosomes, each with two homologs. Both chromosomes have a map length of [0, *R*] where we used *R*=10 such that any real number *r* with O≤r≤ *R* denotes a unique site on the chromosome; the potential number of sites is therefore effectively infinite (e.g. Roze and Rousset 2009). In the first chromosome, mutations are non­neutral, although mutations that are effectively neutral can occur, while in the second chromosome, mutations are fully neutral. The sites on the neutral and non-neutral chromosome are used to calculate a neutral and non-neutral site frequency spectrum, respectively. These are then used as input datasets for the DFE inference software (see below).

Individuals go through a life cycle similar to that of SFS_CODE (Hernandez 2008). Specifically, individuals are sampled with a probability relative to their fitness standardized by the mean fitness of their sex. The fitness of the *i*th individual, *w_i_* is defined as

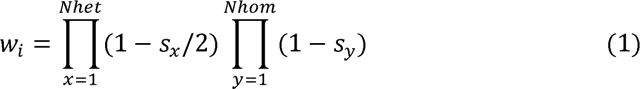

Thus fitness is the product of the effects of all *Nhet* heterozygous mutations and all *Nhom* homozygous mutations, thereby using a model of selection where individuals effects are additive within a locus (i.e., *h*=1/2 for all mutations) and multiplicative between loci. Selection coefficients s are sampled from a gamma distribution (see parameterization below). Parents each provide one of their two homologs to their offspring after a number of recombination events, sampled from a Poisson distribution with mean *R=*10, has occurred. Mutations then occur in offspring and the number of mutations is sampled from a Poisson distribution with a mean of *U*=10 mutations per chromosome per generation, yielding a per genome mutation rate similar to that of some eukaryotes (Haag-Liautard et al. 2007). The number of neutral and non-neutral mutations are sampled independently with the same mean. The selection coefficients of new mutations are sampled from a gamma distribution with a mean of *S=*2*N_e_s* and a shape parameter of *β.* Sampled values of *S* are re-scaled to *s* by calculating *s=S*/2*N_e_* using the estimated *N_e_* for that parameter set (see below). Offspring are created by randomly sampling a mother and a father for each offspring, until *N* viable (w>Q offspring have been produced with a constant 1:1 sex ratio. Once *N* viable offspring have been formed, these will constitute the new parent generation.

To estimate *N_e_* we added a single, neutral, linked locus to the centre of the non­neutral chromosome and mutated the locus at a Poisson-distributed rate with a mean of 1 by adding a random value sampled from *N*(0,l) to the allelic value, thus having the locus satisfy *V_m_* = 1, which is the new genetic variance from mutation appearing per generation (Lynch and Hill 1986; Barton and Turelli 1989; Keightley and Otto 2006). The *N_e_* of the population was defined as the mean (over 50 replicates) steady-state variance at the neutral linked locus after a burn-in period of 10*N* generations. Without selection, the expected variance at this locus is *NV_M_ = N_e_V_M_* (Lynch and Hill 1986) and this was confirmed by simulation (Supporting Information; S1). Mutational effects were sampled as *2N_e_s* (Hernandez 2008), then re-scaled by dividing the sampled value with *2N_e_* before applying it in Eq. 1. To accomplish this, *N_e_* was estimated before the Wright-Fisher simulation by plotting a standard curve of estimated steady-state *N_e_* against an assumed *N_e_* (the value used to re-scale *2N_e_s* to s) for a given gamma distribution of selective effects. This allowed us to know the approximate steady-state *N_e_* which would emerge in our Wright-Fisher simulation before we used these to simulate data for SFS construction (Supporting information; S2).

Data for calculating SFSs were outputted from the simulation after a burn-in period of 10*N* generations, after which the full neutral and selected genomes of 10 random individuals (i.e. 20 haplotypes) where outputted and used to calculated 1 neutral and 1 selected unfolded SFS. Unfolded SFSs were used throughout.

To infer the mean scaled selection coefficients of mutations *S*=2*N_e_s* and the shape parameter *β,* we used polyDFE v1.11 (Tataru et al. 2017; Tataru and Bataillon 2020). To determine whether these parameters could be obtained using our simulation setup, we estimated them from SFSs calculated from simulated populations with the same *S* and *β* combinations that Tataru et al. (2017) used to test the inference of purely deleterious DFEs (see their Fig. 4). We found that polyDFE could accurately infer them from our simulations (Supporting Information; S3) For each single-species polyDFE run, an input file with 50 neutral and 50 selected SFS was used, wherein each pair of neutral and selected SFS was calculated from the basis of a single replicate of a Wright-Fisher simulation (hence 50 Wright-Fisher simulations per polyDFE run set for single-species DFE). The assumed sequence lengths were 20,000 base pairs. For the multi-species polyDFE runs, an input files with 20 neutral and 20 selected SFSs from 3 different species was used (60 SFS). The polyDFE command lines and init-files used are available in Supporting Information; S4.

The concept of *p_lth_* was previously used when fitting a single DFE model to a collection of SFSs obtained from multiple different species (Galtier & Rousselle 2020). To examine the efficacy and validity of *p_lth_* for fitting multi-species DFEs, we simulated various gamma-distributed DFEs in different populations, calculated output SFSs for each population, and inferred DFEs for datasets containing SFSs from multiple different populations. We repeated this analysis with DFEs inferred from genomic data from different species (Chen et al. 2017).

For single species inferences, we first used the model described to simulate populations with *S*=2*N_e_s* in the range of [1000,10000] with increments of *1000* and *β=*04. Each simulation was replicated 50 times (500 replicates in total). For each simulation set, we then inferred *S* and *β* using polyDFE under *β* different *p_lth_* values in the range [0.0,0.9] with increments of 0.1. The parameter *p_lth_* was implemented, following Galtier and Rousselle (2020), by setting an initial *p_lth_* value and scaling each entry of the selected SFS by 1–*p_lth_.* For each *p_lth_*value, 10 polyDFE runs were performed *(1000* polyDFE runs in total) since the performance of polyDFE is affected by the initial *S* value used. For each combination of *p_lth_* value and simulation set, half of the polyDFE runs were run using randomly sampled initial values of the estimated parameters. These values were sampled from a uniform real distribution *U*(1000, 10000) for *S* and *U*/(0.2,0.8) for *β.* For the other half of the polyDFE runs, the true values of *S* and *β* were used as the initial value of the estimated parameters (following Tataru et al. 2017, Supplemental file S1, page 3). To thoroughly test how *p_lth_* affects the overall DFE inference under weaker selection, we also studied the effect of *p_lth_* on *S* and *β* inference given different combinations of *S*= ***50,250,2500*** and β = 0.15, 0.4 0, 0.6 5. For the simulation sets with *S*=2*N_e_s* in the range of [1000,10000] with increments of ***1000*** and *β=*04, we used standard results from probability theory to calculate the true proportion of lethal mutations resulting from these distributions (Appendix A) and compared it to the value of *p_lth_* under which the highest accuracy of inference was found.

We also studied the effects of *p_lth_* on inference of multi-species DFEs from more than one species, and in particular if *β* is fixed between species. This was studied with both arbitrarily defined weakly deleterious DFEs (which were used because they contained hardly any strongly deleterious mutations), and DFEs inferred from genomic data (Chen et al. 2017, 2020). First, we simulated 3 Wright-Fisher populations with different DFEs with the shape parameters *β* and means *S of β =* 0.15, *S_D_ =* 5 *, β =* 0.40, *S_D_* =20 and *β* = 0.65, *S_D_* =40. We then constructed polyDFE input files using 20 neutral and selected SFSs from each population. Using polyDFE, *S* and *β* were estimated from this input file under four different *p_lth_* values using similar procedure of replication and definition of initial values as for the single-species case. Second, using DFEs estimated from genomic data, we ran the same analysis by simulating the DFEs estimated for populations of *Arabidcpsis ¦yrata, Capsella grandflora* and *Zea mays* using the data of Chen et al. (2017). We compared the likelihood of the estimated parameters under 4 different *p_lth_* values, so that they cover cases when the initial values were correctly defined and when these were purposely highly inaccurate. Following Galtier & Rousselle (2020), we also studied the effect of *p_lth_* on *S* inference from these DFEs given different fixed values of *β* (as opposed to jointly estimated). Confidence intervals of estimates shown in plots were calculated as 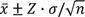 with 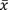 being the mean value, *n* being the sample size, and *Z* being the 97.5 percentile value of the *t*-distribution given *n*–1 degrees of freedom.

## Results

### Analytical results

To illustrate the utility of including a *p*_lth_ parameter in DFE inference and determine where it might be most useful, we first present analytical results demonstrating which deleterious alleles are unlikely to be present in a population sample. Assume that mutations are introduced at a rate θ =4*N_e_µL* for *N_e_* the effective population size, *µ* the per-site mutation probability per generation, and *L* the number of sites where selected sites are likely to arise (e.g., nonsynonymous sites;(Galtier 2016)). A focal mutation has a scaled selection coefficient *S = 2N_e_s* as a heterozygote, and is in the population at a frequency *x.* If we sample *n* haploid genomes, then the expected probability that this mutation is present in *i* of *n* samples is (Tataru et al. 2017, Equation 1):

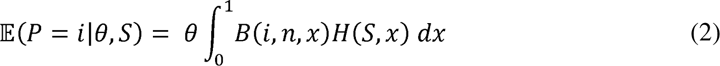

Where *B* indicates the binomial distribution of sampling *i* selected alleles from *n* genomes, and *H* the expected time that the allele spends between frequencies *x* and *x+S x* (Wright 1938):

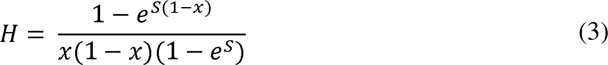

Note there are a few differences between Eq. 3 and the version in Tataru *et al*. (2017). Eq. 2 scales *s* by *2N_e_* whereas Tataru *et al*. (2017) use *4N_e_* scaling, due to the different manner in which heterozygote fitness is calculated. That is, we consider mutations to be additive with a dominance coefficient of *h=*1/2, such that the fitness effect of a single heterozygous and homozygous mutation is 1–*s*, and 1–*s*, respectively. *S* is also multiplied by −1 in Eq. 3, so a positive value denotes a deleterious mutation with that selection magnitude.

Using Poisson Random Field theory (Sawyer and Hartl 1992; Sethupathy and Hannenhalli 2008), the probability of observing a certain number of sites with *i* selected alleles is Poisson distributed with mean 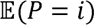 (Eq. 2). Hence, the probability that no sites will have *i* selected alleles is 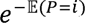, and the probability that no selected alleles are present at all equals 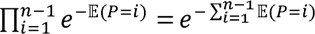. The repository with the code used for all simulations also contains a *Mathematica* notebook to perform these calculations (see Data Availability).

Fig. 1 outlines the probability that no selected allele will be present in a sample of genomes, for different sample sizes and mutation rates. We see that if the mutation rate is low (*θ* = 0.1) then even weakly-deleterious mutations (*S >* 10) are unlikely to be present. For *θ =* 1, while a small sample is also unlikely to contain strongly deleterious mutations (*S* > 100), a large sample size of 100 is likely to capture some strongly deleterious mutations with 100 < *S* < 1000. A very large sample can capture even more strongly deleterious mutations if the mutation rate is high (*θ* = 10). However, under all cases considered, it becomes unlikely to sample extremely deleterious mutations *(S* approaching 5000), even with large sample sizes and mutation rates. Overall, for realistic sample sizes and mutation rates, once *S* exceeds 100 it becomes difficult to capture strongly deleterious mutations in a population sample, and it is in this parameter space that *p*_lth_ might be useful in capturing the proportion of these deleterious mutations. A *p*_lth_-type parameter might also be necessary to calculate the fraction of all deleterious variants if the population mutation rate is very low.

**Figure 1:**
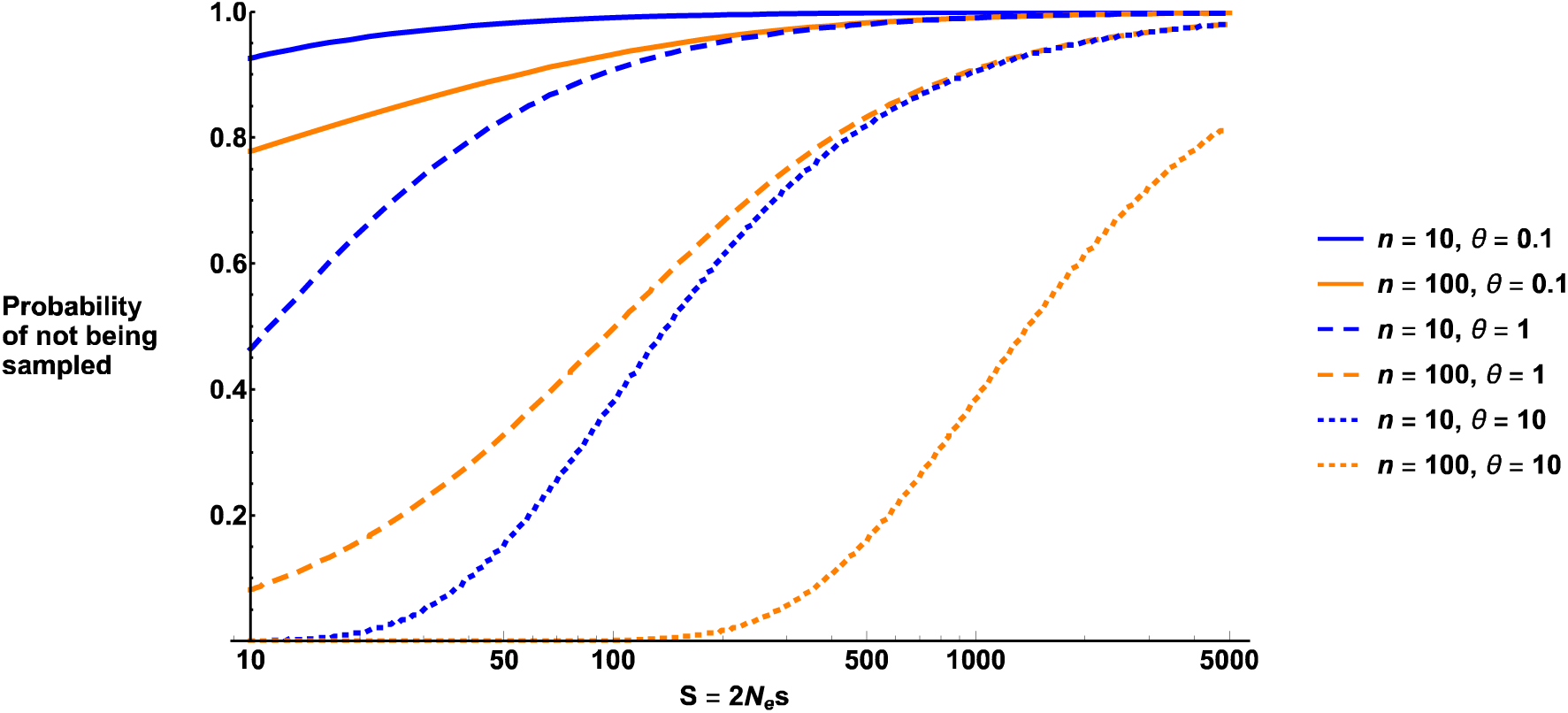
The probability that a deleterious mutation with scaled selection coefficient *S* will not be present in a sample of *n* genomes. Results are plotted for different *n, θ* values as denoted by the legend. Note that we do not plot *S* < 10 since calculations do not assume neutrality and break down as *S* approaches 0. Strongly deleterious mutations are less likely to end up in a sample of haplotypes, making it is difficult to differentiate the effects sizes by analysing site frequency spectra.

### Model-fitting for single-species DFEs

In simulations with a high *S,* the accuracy of inference of *S* and ft was affected by *p_lth_* (Fig. 2A-B) and the accuracy of inference in 4/10 of all cases was highest under a non-zero value of *p_lth_* (Fig-2C-D). In these cases, the accuracy of inference (measured as the difference between ***log***_2_ (*estimate/simulation*) and 0) decreased by a value between 0.13 and 0.99. In terms of the actual estimates of *S,* this means that in the four cases where *p_lth_* improved the accuracy of inference, *S* was estimated 3%-49% (depending on simulation set) more accurately than in the corresponding set where *p_lth_ =*00 . In general, *S* inference becomes more inaccurate as *p_lth_* increases, however the accuracy of inference of ft is less affected unless *p_lth_* is very high (Fig 2). At low mutation rates, high accuracy of inference can be achieved under a high *p_lth_,* although this comes at the expense of the accuracy of inference of β (see Supporting Information; S7).

**Figure 2:**
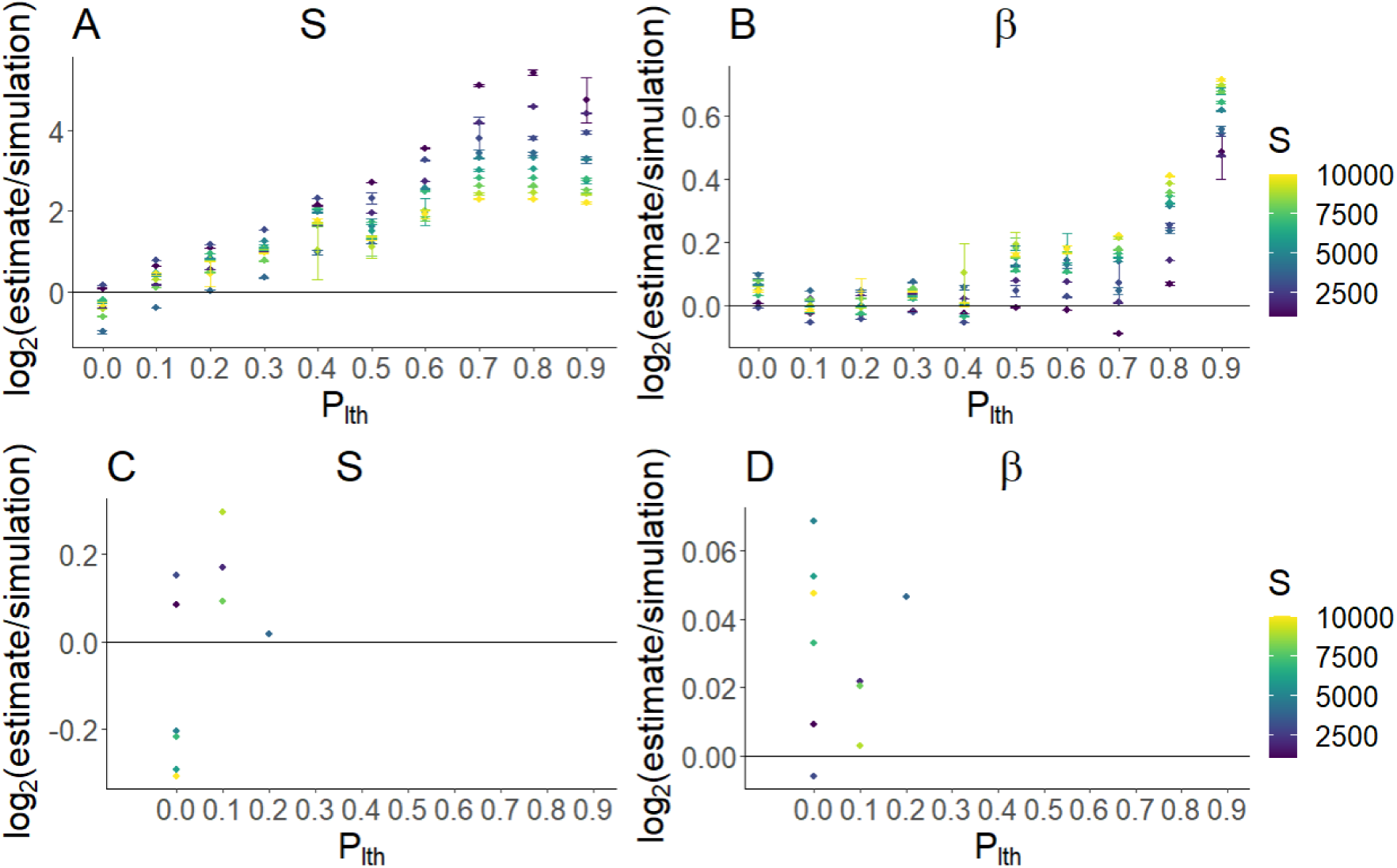
Mean accuracy of inference for *S* (A) and *β* (B) under the assumption of different values of *p_lth_* for different Wright-Fisher simulation sets (differing in the simulation *S,* ranging from 1000 to 10000). Each point shows the mean value of 10 polyDFE replicates. Bars denote 95% confidence intervals (if they are not visible then they lie within the point). For each simulation set (differing in *S;* color), bottom panels show the subset that had the highest accuracy of inference of *S.* For these subsets, the accuracy of inference for *S* (C) and *β* (D) are shown. A value of 0 is equivalent to perfect accuracy of inference. The results show that for each simulation set (these differ in their value of *S*), the best accuracy of inference is found when *p_lth_* is either 0 or very low (*p_lth_ =* 01, *p_lth_ =*02) when a DFE model is fitted to SFS data from a single species.

Under low values of *S* and varying values of *β*, non-zero values of *p_lth_* generally decreased the accuracy of inference, highlighting that *p_lth_* only improves the accuracy of inference when *S* is high (Fig. 3).

**Figure 3:**
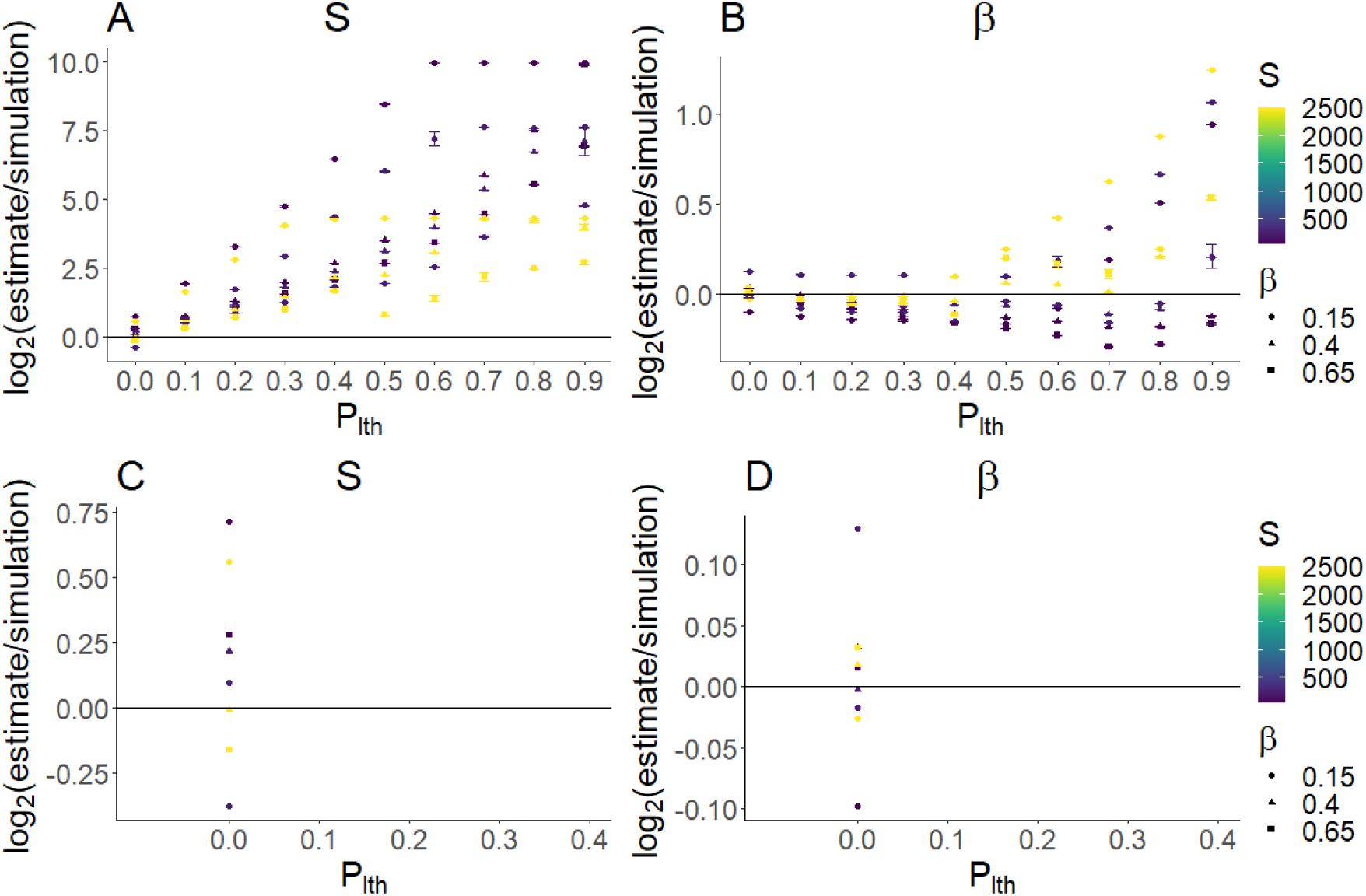
As Figure 2, but instead using 9 simulation sets with different combinations of *S* and *β*(*S* = 50, 250, 2500 and *β* = 0.15,0.40,0.65).

By calculating the true proportion of lethals (Appendix A), which can be used as a lower bound for the proportion of strongly deleterious mutations, we found that the *p_lth_* value resulting in the highest accuracy of inference (Fig. 2, 3) were not equivalent to the true proportion of lethals (Fig. 4), with the true proportions being much higher than those inferred using *p_lth_*.

**Figure 4:**
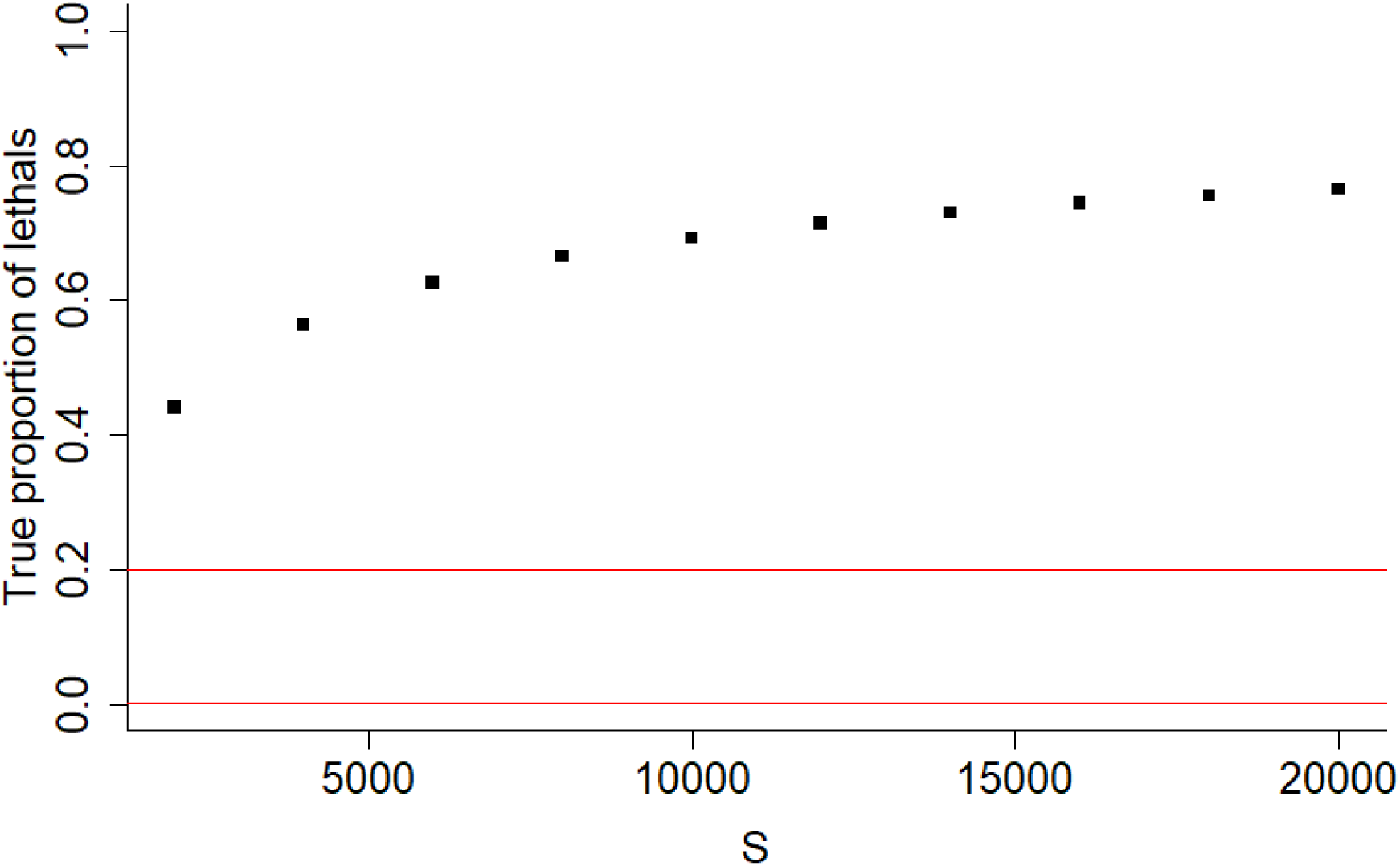
The true proportion of strongly deleterious mutations as a function of *S* (same values as used in Fig. 2) given *β* = 0.40 and *N_e_* = 500 (Appendix A). Red lines show the interval containing *p_lth_* value which resulted in the highest accuracy of inference (Fig. 2, 3). The results show that the lower bounds for true proportion of lethals for each simulation set (points) is higher than the *p_lth_* which results in the best accuracy of inference (0.0-0.2, i.e. between the red lines).

### Model-fitting for multi-species DFEs

We simulated weakly deleterious DFEs for multiple species that had essentially zero probability of yielding lethal mutations (Supporting Information; S5). Despite this setup, we found that the likelihood of the estimated parameters could be artificially maximized when assuming a high *p_lth_* value, when inferring a joint set of DFE parameters from SFSs across different species (Fig. 5A). When a DFE model is fitted to SFSs from multiple species, the model likelihood is artificially maximized because a non-zero value of *p_lth_* pushes the entries of the different SFSs closer together by making them numerically similar (Fig. 6). That is, species with very different SFSs will seem to have more similar SFSs under a high value of *p_lth_,* because *p_lth_* will result in the entries of these different SFS all being closer to 0. Because of this, fitting a single DFE model to joint SFS data yields a better model likelihood once *p_lth_* is applied. Thus, we found that *p_lth_* can erroneously result in a high likelihood of a model despite essentially no strongly deleterious mutations being present.

**Figure 5:**
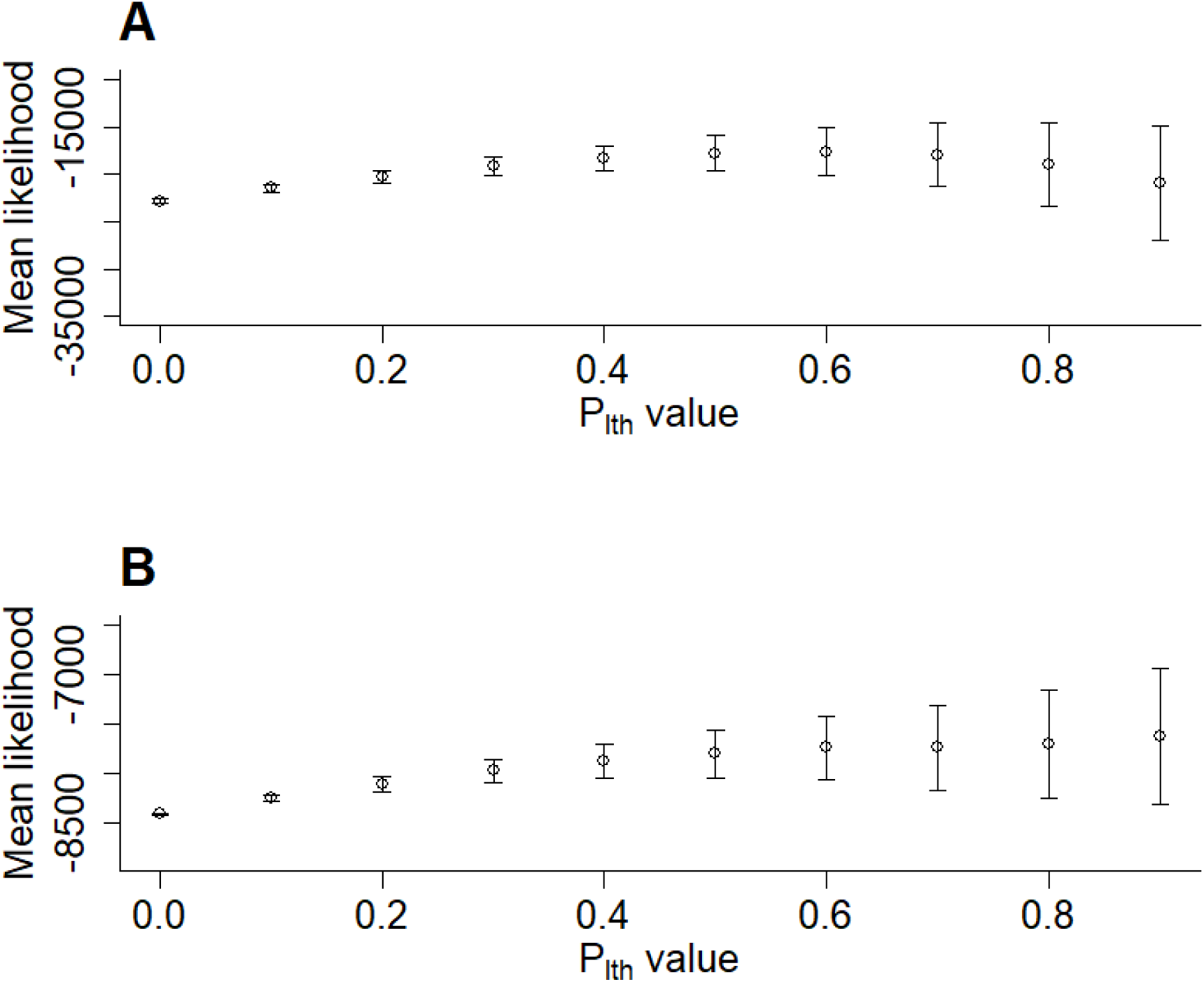
Mean likelihood with 95% confidence intervals returned by polyDFE for a multi­species DFE model against 10 *p_lth_* values. *β* was inferred and *S* was assumed fixed at a randomly sampled value. polyDFE runs were conducted on 20 neutral and selected SFSs from 3 different simulated DFEs. **(A)** Model likelihood for inference based on simulated weakly deleterious DFEs with the shape parameters ft and means *S* of *β* = 0.15, *S* = 5, ft = 0.40, *S* = 20 and *β* = 0.65, *S* = 40 combined into one data input file. **(B)** Model likelihood for inference based on the DFE of three populations simulated parameterized with data from Chen et al. 2017 using DFEs inferred for *A. lyrata, C. grandiflora* and *Z. mays.* The results show that (A) model likelihood can be maximized under a high value of *p_lth_* despite little or no strongly deleterious mutations being present in the DFE because *p_lth_* makes different SFSs more similar and (B) this effect can also occur for DFE estimated from natural populations.

**Figure 6:**
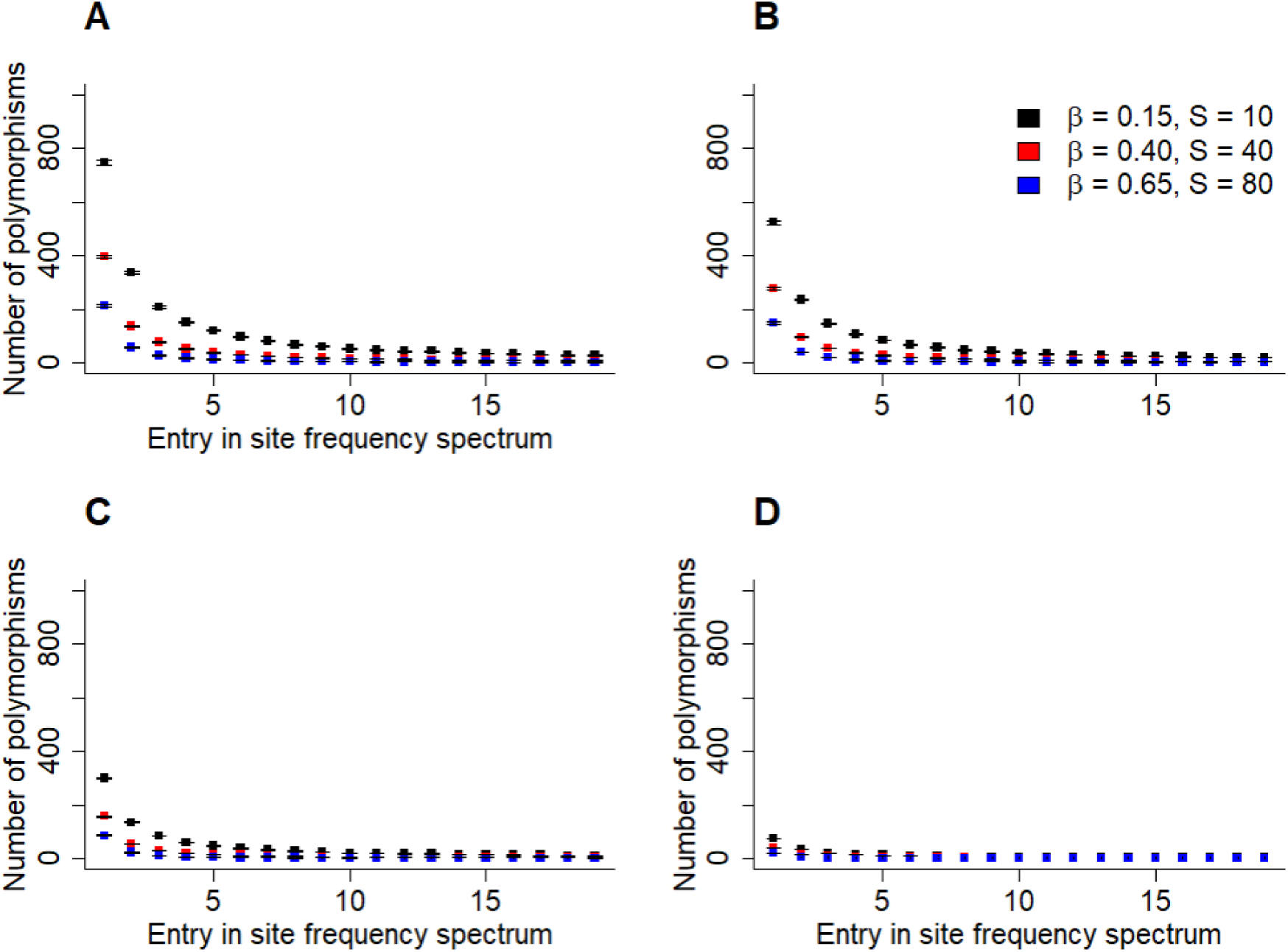
Value of the 20 entries of the SFSs of 3 different simulated DFEs (black, red, blue) under four different *p_lth_* values: (A) *p_lth_ =*00, (B) *p_lth_ =*05, (C) *p_lth_ =*06, (D) *p_lth_ =* O.7.Confidence intervals fall within points as shown. The results show that increasing the *p_lth_* value pushes the entries of different SFS together by making them more numerically similar.

*p_lth_* can also artificially maximise the likelihood of DFE models when simulating DFEs estimated from genomic data (Fig. 5B). Whether the likelihood of the DFE model levels off or keeps increasing with *p_lth_* depends on the sampled initial values for the polyDFE runs. For high *p_lth_* values, the initial values sampled for each polyDFE run becomes increasingly important. This seems to happen as the likelihood of the inferred DFE model drops substantially if the initial *S* fed into polyDFE was low, and the SFSs are modified by a high *p_lth_* value resulting in a seemingly very deleterious selected SFS. Because of this effect, 95% confidence intervals on the mean likelihood increase as *p_lth_* increases (Fig. 5). The parameter *p_lth_* can also artificially maximize the likelihood of a DFE model in cases where *β* is fixed and *S* is inferred, similar to the approach used by Galtier & Rousselle (2020) (Fig. 7).

**Figure 7:**
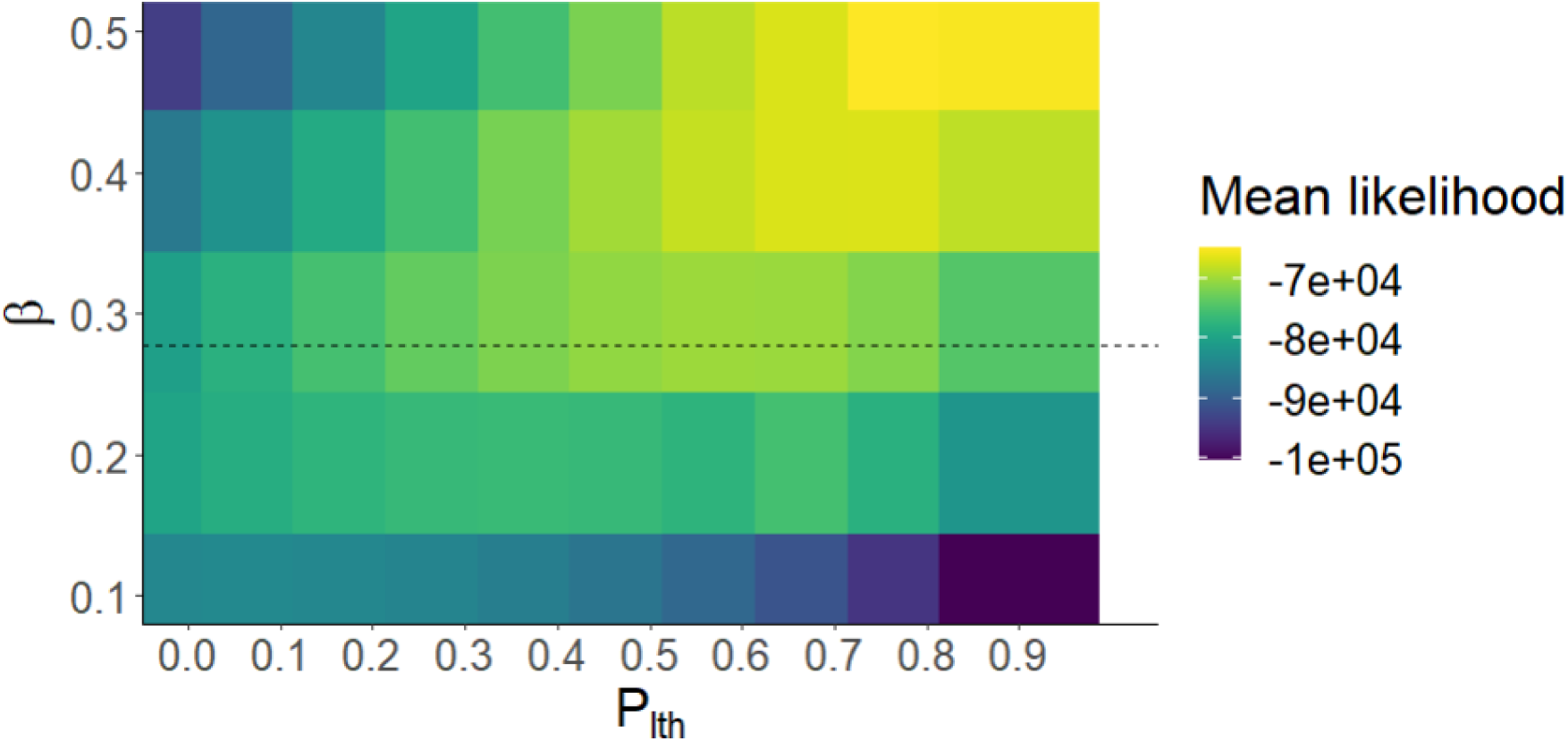
Mean likelihood returned by polyDFE for estimates of *S* given combinations of fixed *β* and *p_lth_.* A multi-species DFE where 20 neutral and selected SFSs from DFEs based on those estimated for *A. lyrata, C. grandiflora* and *Z. mays* by Chen et al. (2017) were combined was used a data input for polyDFE. The dashed line show the mean ft estimated for *A. lyrata, C. grandiflora* and *Z. mays.* The results again shows that model likelihood can be artificially maximized under the assumption of a high value of *p_lth_* and the extent to which this happens depends on the initial values supplies to the DFE inference software.

### Exploring wider parameter space

The effect of *p_lth_* on the accuracy of inference was also tested under larger population sizes and a larger number of sampled haplotypes (Supporting information; S6). While *p_lth_* was less useful for correcting the underestimation of the mean *S* when a larger population size was used, the results remained qualitatively similar such that *p_lth_* had a more positive effect on the accuracy of inference as *S* was increased. The ability of *p_lth_* to artificially maximize the likelihood of a model fitted to SFS data from multiple species was also tested under larger simulated population and sample size and this yielded a result that was qualitatively the same (Supplementary Information; S6). That is, *p_lth_* will also artificially maximize the likelihood of a DFE model based on multiple species when these species have larger populations. This makes intuitive sense given the mechanism through which *p_lth_* seems to cause the artificial maximization of likelihood; that is, the entries of different SFSs will become numerically similar as a higher *p_lth_* is used, regardless of the sample size.

We also investigated inference in a single species using a five-fold lower mutation rate. Here, higher values of *p_lth_* can result in the best accuracy of inference for *S,* however, this happens at the expense of poor accuracy of inference of β (Supporting information S7).

## Discussion

Improving the accuracy with which we can infer the DFE is important since this is a fundamental parameter of evolutionary genetics with implications for many branches of biology. Constructing an accurate description of the DFE boils down to constructing a model which can describe a wide range of effect sizes of mutations at a broad range of frequencies, and hence can be applied to many different species. This is not a trivial problem, because while several models consisting of parametric distributions have been suggested, including gamma, exponential and lognormal distributions, the evidence shows the DFE of deleterious mutations follows some complex and multi-modal distribution rather than a simple parametric one (Nielsen and Yang 2003; Sawyer et al. 2003; Eyre-Walker and Keightley 2007; Kousathanas and Keightley 2013). Thus, fitting a single parametric distribution model to a deleterious DFE can result in a biased estimate, and this bias is especially prone to affect the inferred proportion of strongly deleterious mutations (Kim et al. 2017). Our results illustrate that once strongly deleterious mutations are present, expanding the typical gamma distribution model of a DFE to one which accounts for a proportion of strongly deleterious mutations (Galtier and Rousselle 2020) can in principle improve the accuracy of inference, although it can artificially increase the likelihood of the inferred parameters when a single DFE is fitted to data from different species or populations.

We show that when inferring a DFE from a single species, a model under which *p_lth_* >0, can sometimes yield the highest likelihood even when the DFE does not contain a fraction of *p_lth_* strongly deleterious mutations. This result highlights that *p_lth_* is not the true proportion of lethals and should not be viewed as such. Future developments might produce a more useful correcting factor that accounts for underrepresentation of strongly deleterious mutations while being compatible with classic parametric distributions such as gamma distributions. This would likely involve first calculating how much underestimating might occur, assuming some gamma distributed DFE, and part of answering this would be to calculate the mean effect size of mutations with 2*N_e_s>*2*N_e_*.

Further, the classic notion of a model likelihood is not well-defined when using *p_lth_,* since *p_lth_* is used to multiply all entries of the SFSs by 1–*p_lth_.* This rescaling results in the data itself being modified and consequently two DFE models with differing values of *p_lth_* will be models on different datasets. This means that the likelihood of two different DFE models with different *p_lth_* values cannot be compared in the same way. This explanation does not mean that there is a problem with the concept of model likelihood in general; rather, it instead means that model likelihood cannot be used for model selection in the same way when *p_lth_* is used, since different *p_lth_* models are effectively attempting to explain different datasets.

Given SFS data from a single species, our results illustrate that DFE inference performs better on strongly deleterious DFEs when entries in the input SFS are modified by some value *p_lth_* to reflect the proportion of strongly deleterious mutations that will not end up in a haplotype sample. This is in line with the results of Fig. 4A in Tataru et al. (2017) showing that mutational effects are underestimated when the input DFE is strongly deleterious, indicating that strongly selected mutations are not fully detected or represented in the haplotype sample. This is one parameter space where including *p_lth_* may help with improving the accuracy of inference, albeit the resulting *p_lth_* values are unlikely to reflect the actual proportion of strongly deleterious mutations. This means that although including some value *p_lth_* may improve the model likelihood, *p_lth_* cannot reliably be used to infer the proportion of strongly deleterious mutations. A recent study wherein a DFE model was derived on completely different principles (by considering gene regulatory networks and metabolic pathways) also concluded that no parametric distribution suffices to describe the DFE of strongly deleterious mutations, and that a satisfactory model must involve some extra class of strongly deleterious mutation akin to *p_lth_* (Brajesh et al. 2019). These findings suggest our conclusion has broad generality beyond the Wright-Fisher model. In practice, information about the optimal value of *p_lth_* to use when fitting a DFE model to a particular species may eventually be determined *a priori* based on the species in question. For example, an application of Fisher’s geometric model (Fisher 1930) to DFE theory yields the prediction that as the complexity of an organism increases, a larger proportion of the DFE should be strongly deleterious (Lourenço et al. 2011; Tenaillon 2014). This has been tested several times and found to be consistent with data by using the level of pleiotropy as a proxy for the complexity of an organism (Martin and Lenormand 2006; Huber et al. 2017).

It is well-known that the likelihood of an individual DFE model drops when fitted to multiple species (Huber et al. 2017; Galtier and Rousselle 2020), which highlights that the DFE differs among species (Eyre-Walker 2002; Lynch and Conery 2003). In this study, we also show that *p_lth_* can artificially improve the likelihood of parameter estimates when a single DFE is fitted to multiple species. This result has several important implications. First, it illustrates the point that while the data strongly suggests the DFE of deleterious mutations usually follows a gamma and lethal model, it can still result in a misleading description of the DFE when fitted to data from multiple species. Second, it shows that since *p_lth_* consists of a modification of the data, we should bear this in mind when assessing the likelihood of a model; we find the likelihood of a DFE model is maximized under a high *p_lth_* in cases where the true DFE is known to be very weakly deleterious and known to not contain a proportion of *p_lth_* strongly deleterious mutations. Because of this, we can conclude that if the likelihood of a given DFE model for multiple species is maximized under the assumption of some *p_lth_* >0, this should not be considered evidence that the DFEs being modeled does indeed contain a proportion of *p_lth_* strongly deleterious mutations. Third, our results offer an explanation of the mechanism by which this effect occurs, namely that *p_lth_* makes entries of different SFSs more numerically similar, thereby inflating the likelihood of the resulting DFE model. Several studies model DFEs as following a gamma and lethal model, or gamma and “point mass” model (Eyre-Walker et al. 2006; Elyashiv et al. 2010), but when such models involve applying a correction factor to SFSs, they seem unsuitable for fitting a single DFE to data from a group of species.

While our study is limited to purely deleterious DFEs, more work on improving the accuracy of inference for DFEs with beneficial mutations is also needed, since recent work shows that the occurrence of strongly beneficial mutations (much like strongly deleterious mutations) can make DFE inference inaccurate (Booker 2020). Even in the study of purely deleterious DFEs, attention has almost exclusively been focused on the effects of SNPs (resulting from point mutations), but the DFE of INDELs (insertions and deletions) is understudied and accounting for such mutations might require a slightly different modelling approach (Barton and Zeng 2018). As in our study, mutational effects are most often assumed to be additive, however the average dominance coefficient of new mutations appears to be substantially lower than *h*=1/2 (Mukai et al. 1972; Simmons and Crow 1977; Lynch et al. 1999; Fernández et al. 2004; Spigler et al. 2017). While it is implicitly assumed that current DFE inference algorithms remains accurate when the assumption of additive dominance is violated, very little work has been done to test this assumption (but see Wade et al. 2022). Similarly, other “gamma + lethal” DFE models exist and since these differ slightly from *p_lth_* implemented in Rousselle & Galtier (2020), testing their ability compared to the *p_lth_* implementation of a “gamma + lethal” DFE model is a topic worthy of further research (Boyko et al. 2008; Kim et al. 2017).

Experimental evidence suggest that the DFE of deleterious mutations follow a bimodal or perhaps even multi-modal distribution, meaning that a good model fit may not be possible under classic parametric distributions such as exponential, gamma, or lognormal (Nielsen and Yang 2003; Sawyer et al. 2003; Eyre-Walker and Keightley 2007; Kousathanas and Keightley 2013). Because of this, it would be worthwhile for future research to explore whether DFE model fitting under different distribution could result in an improve model fit. In a recent study, the DFE was represented by using a non-parametric distribution in the form of several non-overlaping uniform distributions (Johri et al. 2020). Some of the current DFE inference software can also be set to infer a DFE where selection coefficients take discrete values rather than necessarily conforming to a single continuous distribution (Tataru et al. 2017). Expanding the set of distributions typically used for DFE model could prove to be fertile grounds for new research and result in more accurate models.

## Conclusion

While the parameter *p_lth_* can in principle improve the accuracy of inference, obtaining a good model fit under some non-zero *p_lth_* value should not be viewed as evidence for a proportion of *p_lth_* mutations segregating in the population in question. This is because it can be shown that *p_lth_* is not equivalent to the true proportion of lethals. When inferring a single DFE for a group of species or populations, the usage of *p_lth_* is also problematic since it modifies the data, resulting in different SFSs becoming more alike and artificially increases the likelihood of the model inferred. Thus, comparing the likelihood of two models with different *p_lth_* values cannot be done in a standard way, since these two models will effectively have different data. We have presented a detailed study of some of the problems with *p_lth_* as a concept which will be useful to anyone modelling the DFE, especially with regards to avoiding model artifacts.

## Data availability

The simulation software was implemented in C++ and the full source code is available at https://github.com/r02ap19/DFE_Wright-Fisher01.

## Supporting information

Supporting informaton for main text

## Acknowledgements

We thank Adam Eyre-Walker, Brian Charlesworth, Thomas Bataillon, Jane M. Reid, Roslyn Henry, and Max Tschol for many helpful ideas, comments, and suggestions. We also thank Nicolas Galtier for reading an earlier draft and providing many useful comments and much valuable feedback. Anders P. Charmouh was supported by the University of Aberdeen. Greta Bocedi was supported by a Royal Society University Research Fellowship (UF160614). Matthew Hartfield is supported by a NERC Independent Research Fellowship (NE/R015686/1) and a UKRI Frontier Research Guarantee Grant (EP/X027570/1).

## Appendix A

Here, we show how to calculate the fraction of mutations with s>l given some *N_e_* and some gamma distributed input distribution (Fig. 4). The calculations can in principle be performed for any input distribution where the probability density function and its parameters are known. As an example, we will use DelHSD from Tataru et al. (2017): a gamma distribution with a shape parameter of *β*=04 and *S* = 10000.

For any gamma distribution with shape parameter *β* and scale parameter *θ,* the probability density function is

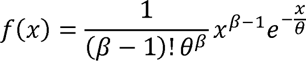

which we can use to find the fraction of mutations with *s*>l by integrating *f* over [0,4*N_e_*]

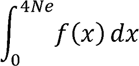

Since *E*[*f*(*x*)] *= βθ,* the probability density function for DelHSD becomes

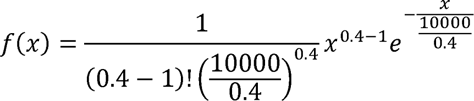

And with *N_e_* = 500, we get

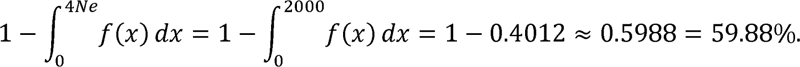

