## Supporting informaton for main text for "Inferring the distributions of fitness effects and proportions of strongly deleterious mutations"

**Supplementary Information**

**S1. Test of** $\boldsymbol{N}\boldsymbol{V}_{\boldsymbol{M}}\boldsymbol{=}\boldsymbol{N}_{\boldsymbol{e}}\boldsymbol{V}_{\boldsymbol{m}}$ **under neutral conditions….....………………………………….2**

**S2. Calibration of** $\boldsymbol{N}_{\boldsymbol{e}}$ **for scaling** $\boldsymbol{2}\boldsymbol{N}_{\boldsymbol{e}}\boldsymbol{s}$ **to** $\boldsymbol{s}$**……………………………………………...…2**

**S3. Test of the accuracy of inference under cases with** $\boldsymbol{p}_{\boldsymbol{lth}}\boldsymbol{=0}$**………...…...……..……..4**

**S4. Command lines and init_files for polyDFE runs……………………………………….5**

**S5. Probability distributions of weakly deleterious DFEs for multi-species tests…...…...5**

**S6. Replication of main results under larger population and sample size………...……...6**

**S7. Additional simulation under lower deleterious mutation rate………………...……...8**

**References………………………………………………………………………………...…10**

**S1. Test of** $\boldsymbol{N}\boldsymbol{V}_{\boldsymbol{M}}\boldsymbol{=}\boldsymbol{N}_{\boldsymbol{e}}\boldsymbol{V}_{\boldsymbol{m}}$ **under neutral conditions**

To check that our model behaves as we would expect under a Wright-Fisher population, we measure if the steady-state variance (after the burn-in period) at the neutral linked locus is equal to $N$ once all mutations are neutral (Lynch & Hill 1986; Keightley & Otto 2006). Fig. S1 shows an example from a simulation replicate, highlighting how $NV_{m}=N_{e}V_{m}.$

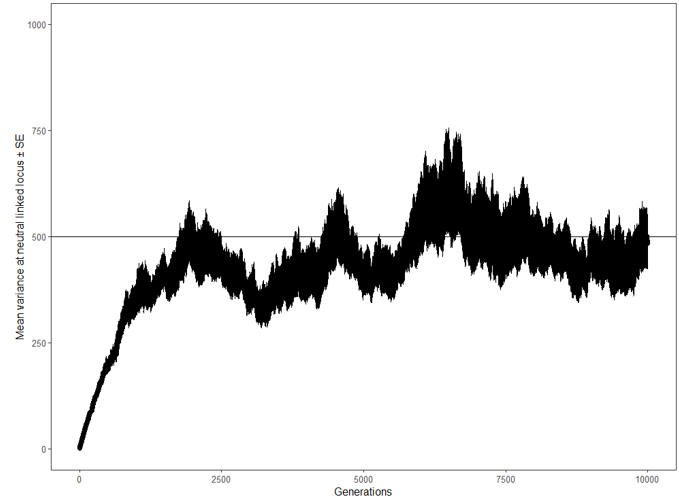

Figure S1: Mean $N_{e}$ of 50 replicates over time with only neutral mutations $\pm$SE. The mean $N_{e}$ after a burn-in period of $10N$ $(5,000$ generations) was $485\pm38$. The straight black line denotes $N=N_{e}$.

**S2. Calibration of** $\boldsymbol{N}_{\boldsymbol{e}}$ **for scaling** $\boldsymbol{2}\boldsymbol{N}_{\boldsymbol{e}}\boldsymbol{s}$ **to** $\boldsymbol{s}$

As shown in S1, $N_{e}$ was estimated by measuring variance at a neutral linked locus (Hill & Lynch 1986; Keightley & Otto 2006). Since $N_{e}$ is used to scale mutational effects sampled as population-scaled mutational effect ($S=2N_{e}s$) to selection coefficients ($s)$, it is necessary to know approximately which value of $N_{e}$ will emerge in a simulation. Because the emerging value of $N_{e}$ is itself affected by the simulation $S$, we estimated an approximate emerging $N_{e}$ by plotting a standard curve of estimated steady-state $N_{e}$ against an assumed $N_{e}$ (the value used to re-scale 2$N_{e}s$ to $s$) for a given gamma distribution of selective effects. This allowed us to know the approximate steady-state $N_{e}$ which would emerge in our Wright-Fisher simulation before we used these values to simulate data for SFS construction.

For each standard curve, 10 x 10 replicate simulations with differing assumed $N_{e}$ (calibrating $N_{e}$) were simulated and the emerging mean $N_{e}$ was calculated (using variance at the neutral linked locus, see S1) and used for re-scaling mutational effects in simulations from which SFSs where calculated (for an example of a standard curve, see Fig. S2). The $N_{e}$ used to scale the mutational effects for the actual simulations was selected as the mean of the emerging $N_{e}$ of the calibration simulations. The $N_{e}$ emerging in simulations from which SFSs where calculated was similar to the approximate $N_{e}$ estimated by this calibration procedure.

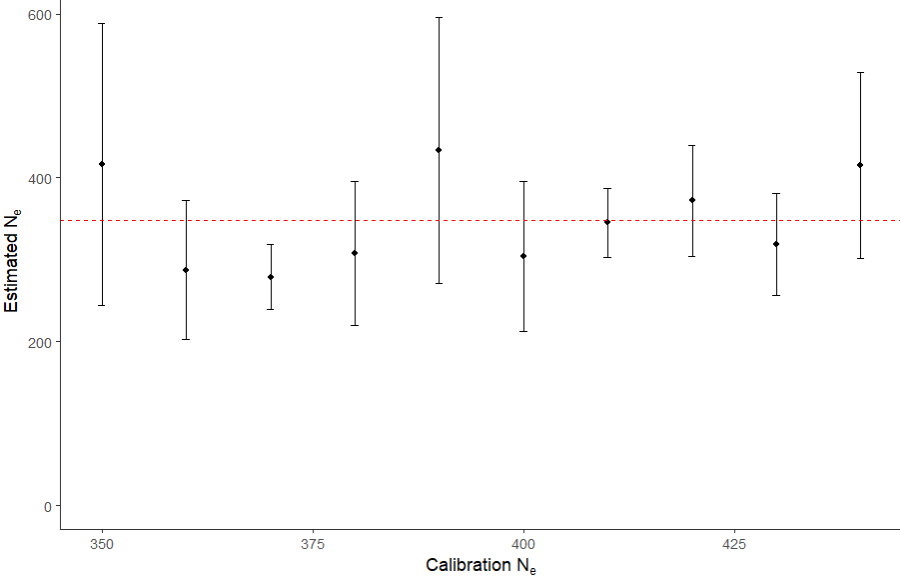

Figure S2: Example of standard curve used to estimate which $N_{e}$ would emerge in a simulation given some $N_{e}$ used for scaling mutational effects and $S$. Estimated $N_{e}$ values as a function of assumed $N_{e}$ values (calibration $N_{e}$) used to re-scaled mutational effects from $2N_{e}s$ to $s$. Each point is the mean of 10 simulation replicates with standard error. For these simulations, $N=500, S=1000.$ For all simulation sets, the mean emerging $N_{e}$ was calculated (dashed red line) and this was used to scale mutational effects in the simulations from which SFSs were calculated.

**S3. Test of the accuracy of inference under test cases with** $\boldsymbol{p}_{\boldsymbol{lth}}\boldsymbol{=0}$**.**

Compared to SFS_CODE (Hernandez 2008), our model implements the idea of $N_{e}$ and recombination slightly differently (see main text for details). To ensure that polyDFE can still accurately infer DFEs from our output, we replicated Fig. 4A of Tataru et al. 2017 to test the accuracy of inference under standard cases with $p_{lth}=0$ (Fig. S3).

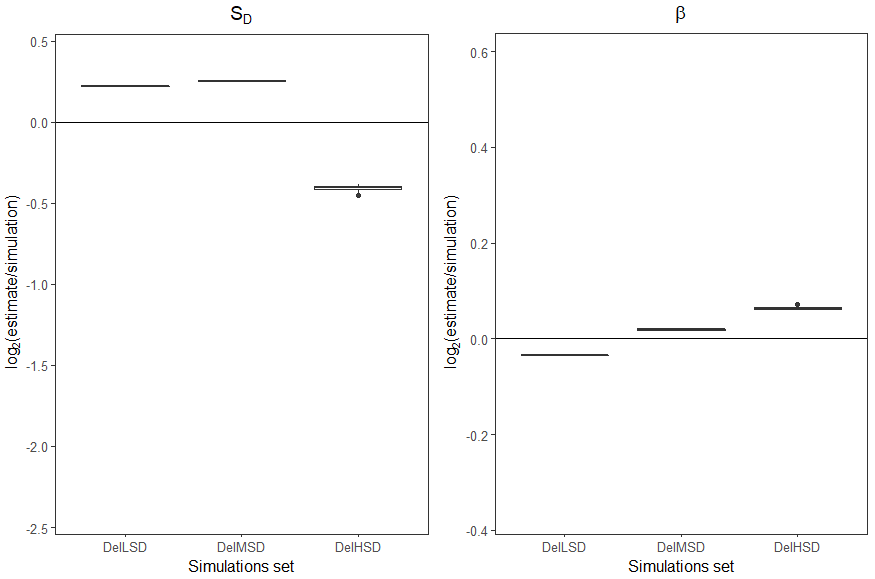

Figure S3: Boxplots of 10 polyDFE runs showing the accuracy of inference of DFE for the gamma distributions DelLSD, DelMSD and DelHSD of Tataru et al. 2017. The left panel shows the accuracy of inference of $S$, while the right panel shows the same for $\beta$. As expected, the results show slightly lower but comparable accuracy of inference to Tataru et al. 2017 despite a slightly different model.

For $S$, the inferred means for the three groups DelLSD, DelMSD and DelHSD were 11.62, 476.77 and 7531.32 and the true values were 10, 400 and 10000. In all cases, the true value of β was 0.40. The remaining discrepancy between true and inferred values under our model likely stems from the fact that SFS_CODE (which is what Tataru et al used) treats $N_{e}$ as a parameter (chosen by the user) and scales mutation and recombination rates accordingly, whereas $N_{e}$ in our model just emerges given the chosen $N$ (see also S2).

**S4. Command lines and init_files for polyDFE runs**

Below are examples showing a polyDFE command line and a polyDFE init file used for the analysis resulting in Fig. 1 in the main text. For details on polyDFE command line and init files, please see the polyDFE documentation; Tataru et al. 2017.

polyDFE was run using the following command line:

./polyDFE -d 0_SFS.txt -i init_model_BandH.txt 1 -v 100 -w > output

For the init files, which contains the following parameters

#ID	eps an	eps cont lambda	theta bar a	S d	b	p b	S b	r (at least #groups-1 of them)

the following line was used

1	1 0.00	1 0.00	0 0.005	0 0.001	1 -1	0 -17694.63 0 0.22	1 0	1 0	0 1 1 1 1 1 1 1 1 1 1 1 1 1 1 1 1 1 1 1 1 1 1 1 1 1 1 1

where -17694.63 and 0.22 in this case are examples of randomly sampled initial values for $S$ and $\beta$, respectively. For more details on random sampling of initial values, please see the main text.

**S5. Probability distributions of weakly deleterious DFEs for multi-species tests**

To test whether high $p_{lth}$ could maximize the likelihood of a DFE in model even in cases where essentially zero strongly deleterious mutations are present, we defined three weakly deleterious gamma-distributed DFEs with different shape parameters $(\beta)$ and means $(S)$. We then fitted DFE models to SFSs from all three distributions (a multi-species DFE model) and studied how the likelihood of the model was affected by changing $p_{lth}$ (see main text). The probability distribution of mutational effects from these three DFEs are visualized in the following (Fig. S4).

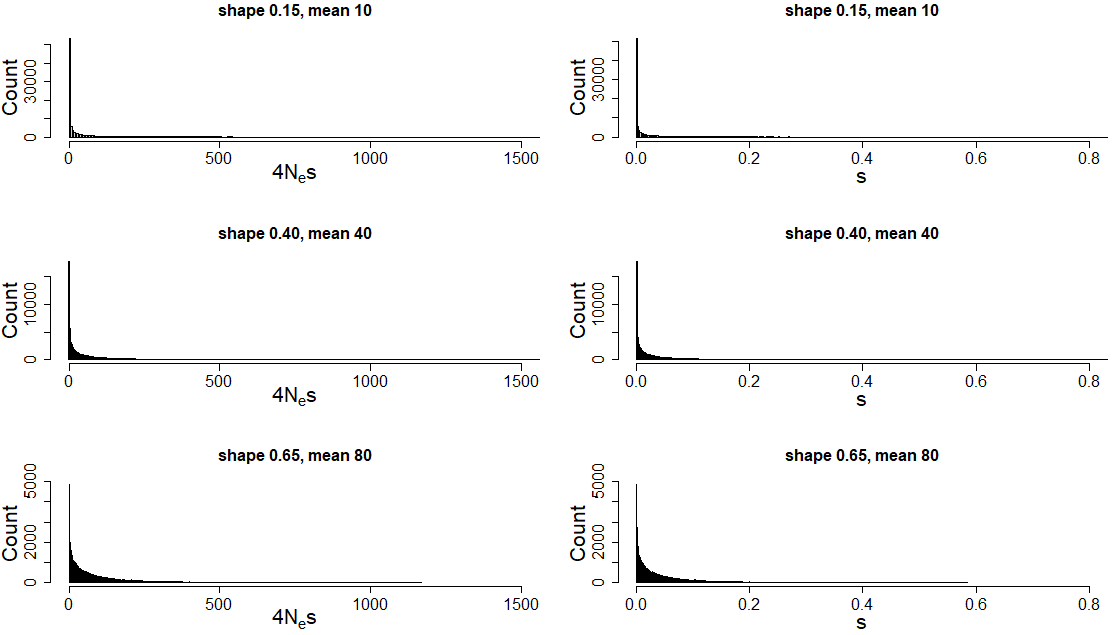
Figure S4: Visualisation of the probability density of 3 weakly deleterious DFEs with the shape parameters $\beta$ and means $S$ of $\beta=0.15, S=10$, $\beta=0.40, S=40$ and $\beta=0.65,S=80$ for simulated Wright-Fisher populations with $N=500$. Mutational effects are shown as $4N_{e}s$ (left column) and $s$-coefficients (right column).

**S6. Replication of the main results under larger population and sample size**

To show examples of the main results under larger population size and larger number of sampled haplotypes, we ran Wright-Fisher simulations with $N=2,000$ and $n=50$ (see main text for details). We then tested how the accuracy of inference was affect by $p_{lth}$ when a DFE was inferred based on SFSs calculated from a single species (Fig. S5). We also tested whether $p_{lth}$ would still artificially maximize the likelihood given higher $N$ and $n$ when a single DFE was inferred for SFSs from multiple species (Fig. S6).

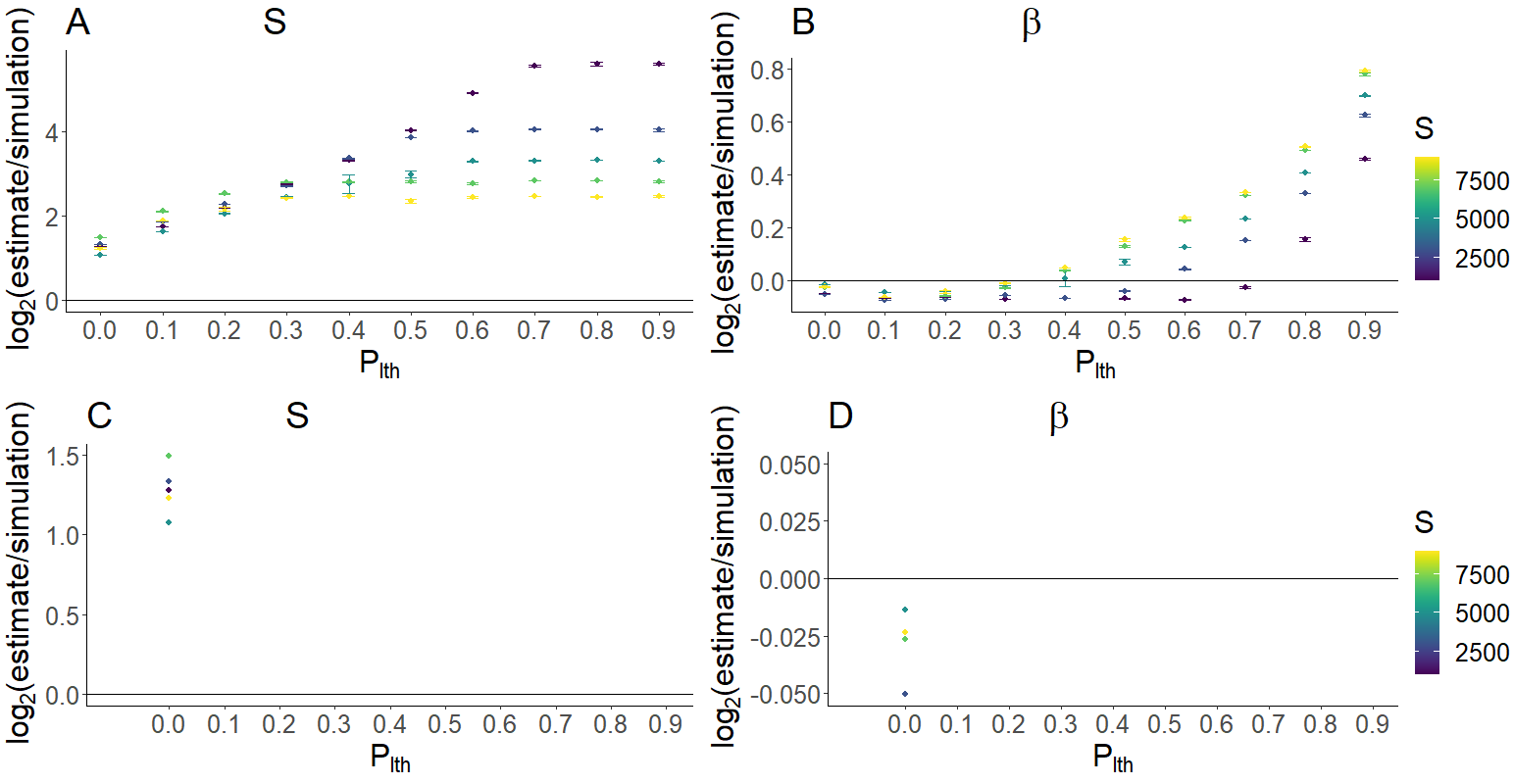
Figure S5: Mean accuracy of inference for $S$ (A) and $\beta$ (B) under the assumption of different values of $p_{lth}$ for different Wright-Fisher simulation sets (differing in the simulation $S$, 2000, 6000, 10000, 14000, 18000). Each point shows the mean value of 10 polyDFE replicates. Bars denote 95% confidence intervals (in most cases, these are not visible as they lie within the point). For each simulation set (color), bottom panels show the subset that had the highest accuracy of inference of $S$. For these subsets, the accuracy of inference for $S$ (C) and $\beta$ (D) are shown.$y=0$ is equivalent to perfect accuracy of inference.

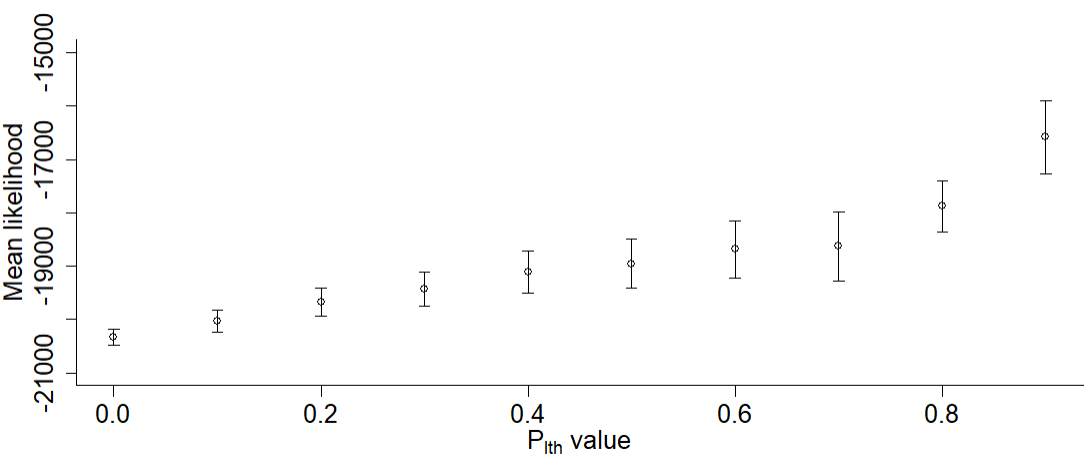

Figure S6: Mean likelihood with 95% confidence intervals returned by polyDFE for a multi-species DFE model against 10 $p_{lth}$ values. $\beta$ was inferred and $S$ was assumed fixed at a randomly sampled value. 20 neutral and selected SFSs from 3 different simulated deleterious DFEs with the shape parameter $\beta=0.40$ and $S$ of $2000, 6000$ and $10000$, respectively, were combined into one data input file. Each point represents the mean value of 10 polyDFE runs.

Under higher population size and higher number of sampled haplotypes, the results remain qualitatively the same: $p_{lth}$ may improve the accuracy of inference if $S$ is high (Fig. S5A). When a DFE is fitted to SFS data from multiple species, high values of $p_{lth}$ artificially maximize the likelihood of the DFE model (Fig. S6) despite no proportion of $p_{lth}$ mutations being present in the dataset (Appendix A in the main text).

**S7. Additional simulation under lower deleterious mutation rate**

To explore how the effect of $p_{lth}$ depends on the deleterious mutation rate, we replicated figure 2 from the main text using a five-fold lower deleterious mutation rate ($U=0.2)$.

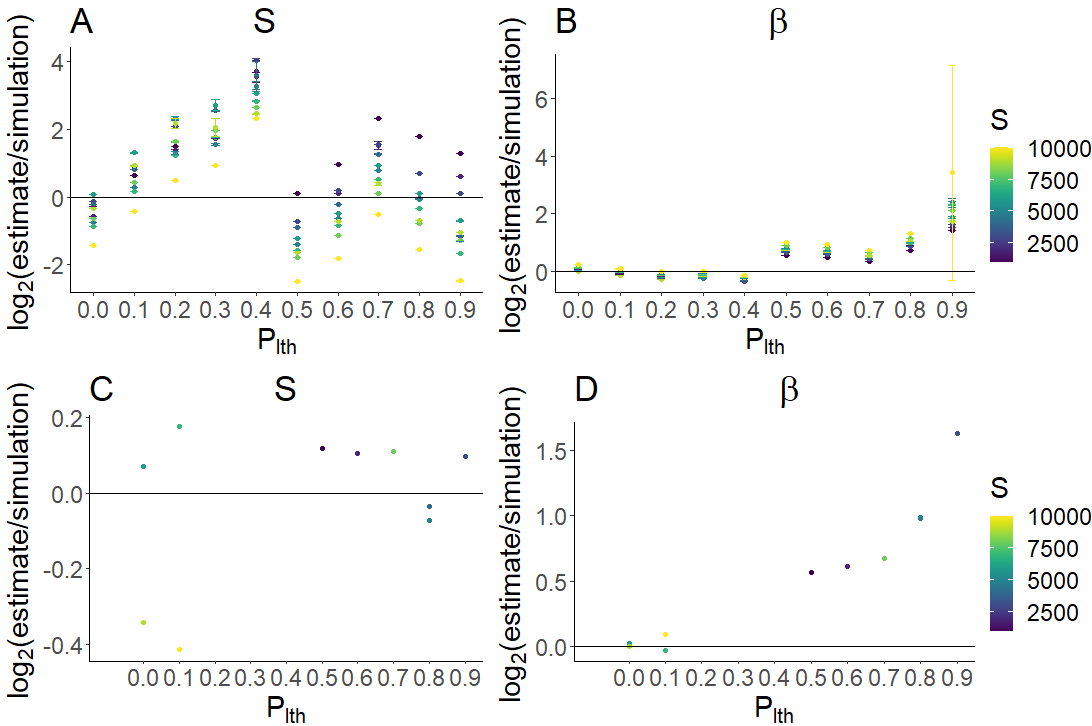

Figure S7: Mean accuracy of inference for $S$ (A) and $\beta$ (B) under the assumption of different values of $p_{lth}$ for different Wright-Fisher simulation sets (differing in the simulation $S$, 2000, 4000, 6000, 8000, 10000, 12000, 14000, 16000, 18000, 20000). Each point shows the mean value of 10 polyDFE replicates (note that V.2 was used here, as opposed to V1.11 for other analyses). Bars denote 95% confidence intervals (in most cases, these are not visible as they lie within the point). For each simulation set (color), bottom panels show the subset that had the highest accuracy of inference of $S$. For these subsets, the accuracy of inference for $S$ (C) and $\beta$ (D) are shown.$y=0$ is equivalent to perfect accuracy of inference.

Unlike single-species simulation under a normal mutation rate, the best accuracy of inference of $S$ can be achieved under a high value of $p_{lth}$, however this comes at the expense of poor accuracy of inference of β. This seems to be because $S$ and $\beta$ are correlated parameters since the mean of a gamma distribution is the product of the shape and scale parameter (Galtier & Rousselle 2020). Once $p_{lth}$ is $\geq0.5$ at low mutation rates, polyDFE seems to favour a DFE model with a lower $S$ that is compensated by a higher $\beta$.
